# Eco-Anthropological factors predicting forest patch use by 3 species of wild Atelid monkeys co-existing with a small-scale farming community in Northeastern Costa Rica, Mesoamérica

**DOI:** 10.1101/2024.02.06.579063

**Authors:** Juan Pablo Perea-Rodríguez, Hugo Carbonero, Rocío Vargas, Claudia Villarreal

**Author notes:** Corresponding Author: Juan Pablo Perea-Rodríguez, Ph.D.

## Abstract

The main conservation risks for wild non-human primates (NHP) in Costa Rica, Mesoamérica, is deforestation and the allocation of lands for agriculture. This is because they result in a mosaic of forest patches that differ in size and ecological properties. NHP, being the vertebrates with the highest risk and rate of extinction, slowly adapt to this rapid environmental change, optimizing their metabolic costs to survive and reproduce. One way to balance these costs is to use forest patches depending on the benefits they provide, such as food, shelter, or social contact. To understand the possible environmental factors that predict the usage of a series of 8 connected forest patches by *Ateles geoffroyi*, *Alouata paliatta*, and *Sapajus imitator* we collected demographic, behavioral, climatological and other environmental data from 2018 until 2021. We used information-theoretic metrics to identify the factors that best explained the presence and behavior of the species of interest in the forest patches studied, and fit the data to a set of models built informed *a priori*. Using the best explanatory factors, we k-fold cross-validated 9 classifier algorithms to identify the best predictive models for the presence of the monkeys studied and their behavioral patterns given the data. Presence was highest in warmer, more humid days, especially when other groups were present in the same patch. Behavioral patterns were different in each patch; monkeys rested more often when other groups of the same species were present, and foraged more during warmer, more humid days, and smaller groups. Predictive models for the presence of the species studied, trained with the 3 best explanatory factors, reached an accuracy between 70-96%, with Gradient Boost Classifier performing the best. In contrast, behavioral patterns were more unpredictable, with the the algorithms tested only reaching between 43-51% accuracy, the AdaBoost Classifier being the best. Our findings suggest that the usage of the 8 forest patches monitored by the monkeys studied depends on patch characteristics, not related to size nor the presence of a reserve, by the presence of other NHP in the patch and the meteorological conditions. Further work on the ecological characteristics of these patches can clarify the mechanisms modulating behavioral patterns.

## Introduction

Wild non-human primates (NHP) are one of the groups of mammals with the highest risk and rate of extinction due environmental change due to human-induced habitat loss and other activities (Estrada et al., 2012, 2013, 2020). As such, NHP adapt to this rapid change balancing their slow development, long lives, high reproductive cost, and sometimes specialized nutritional needs. In areas with high NHP diversity where there are protections set in place, their well-being, welfare, and conservation is dependent on local human activity. This impact, of course, varies in magnitude depending, for example, on the type of agricultural activity and the efforts made to preserve local biodiversity (Estrada et al., 2005). These activities are ultimately linked to human welfare and well-being. Thus, NHP often find themselves in fragmented landscapes with numerous connected patches with different ecological characteristics that influence why they visit them and how often.

Arboreal NHP are especially affected by habitat modification because even some relatively environmentally ‘friendly’ activities can make large areas, of what was originally untouched forest, unusable landscapes to them (Estrada et al., 2005). Thus, in order to understand how communities of NHP respond to the changes in their environment it is important to tease apart the potential effects that different environmental (i.e., abiotic and biotic) factors may have on their behavior and physiology, including those factors relating to cohabitating with humans. This approach can be especially useful in designing data-informed interventions to promote the conservation of wild NHP and their habitats, and to ultimately ameliorate human well-being.

Costa Rica is a relatively small Mesoamérican country, known for its biodiversity, which is estimated to represent 5-6% of the diversity worldwide. There are approximately 8500 species of plants, 220 species of reptiles, 160 species of amphibians, 205 mammals and 850 species of birds. Three of the 4 NHP species found in Costa Rican forests are Atelids (i.e., *Ateles geofroyi*, *Sapajus imitator*, and *Alouatta paliatta*), which relatively larger in size (i.e., 4-7 kg) for Pan American monkeys (Johnson et al., 2023; Zaldivar et al., 2004). The 4th and least abundant species of NHP found in Costa Rica is the Mesoamérican squirrel monkey (*Saimiri oerstedii*), and there has been much success in protecting populations and promoting their reproduction, which has been a decades-long effort (Ceballos et al., 2019). Interestingly, some propose that owl monkeys (*Aotus spp.*) are still present in the in the provinces of Heredia and Limón (Sclater, 1872; Timm, 1988, 1989), but no official census exists of the area for this specific purpose since the late 1800s (e.g., Tafoya et al., 2020; Zaldívar et al., 2004).

Forest fragmentation in Costa Rica has resulted from deforestation going back to several decades of lumber extraction and allocation of lands for agricultural use (Garber et al., 2010). Due to this, the remaining forest is composed of patches that are distant from one another separated by pastures from farmlands, and commercial banana, pineapple, and palm-oil production. These patches often differ in shape, size, and ecological structure and thus provide different benefits and disadvantages to the NHP and other wildlife in the area. NHP show several adaptations to living in fragmented habitats, for example, by changing their activity budgets (Chaves et al., 2011; Gabriel, 2003), diets (Chaves et al., 2023), and social structure (Bolt et al., 2019).

This study had 2 main goals: first, to model the presence/absence of the 3 NHP species (independent of species) in 8 forest patches that varied in size during a 2 year period that included pre- and post-COVID-19 sampling. Second, to model how the three different species of monkeys were using the patches based on their behavior (i.e., resting, traveling, foraging). An information-theory (I-T) approach was used to model the presence/absence and behavior of monkeys found in the patches with 8-9 main effects representing different aspects of their environment, including anthropological, climatological, and demographic factors. We then cross-validated 9 different classifier algorithms trained with the best explanatory factors to build a predictive model. We hypothesized that monkeys would use the different patches monitored due to their ecological characteristics and the possible benefits they may bring, which are dependent on human activity (e.g., agricultural practices).

## Methods

### Study Design

In order to investigate the factors that may explain and predict forest path use of wild Central American spider monkeys (*A. geofroyii*), mantled howler monkeys (*A. palliata*), and white-faced capuchins (*S. imitator*), we mapped 8 forest patches in the study area (4-7 km long) and then monitored them bi-weekly for the presence/absence of the 3 Atelid species of study. Each time we encountered a monkey we recorded their group size, the number of adults and infants in the group, and their behavior (i.e., resting, traveling, foraging) at the time of the encounter.

We sampled all 8 forest patches using the same transects and protocol. We collected data between October of 2018 and December of 2021, unless COVID-19 country wide mobilization restrictions were being implemented. Environmental data included climatology extracted from an online database (i.e., ambient temperature, ambient humidity, rainfall, and wind speed). The goal of the analyses was to model the presence/absence and the behavior of the 3 Atelid species of interest in the 8 patches monitored based on several biotic and abiotic environmental factors, including demographic factors, local climatological patterns, the proximity to reservation area, and the presence of other monkey groups in the patch.

### Study Species

The members of the taxonomic family *Atelidae* include all howler (*Alouatta spp.*), capuchin (*Sapajus spp.*), spider (*Ateles spp.*), and wooly monkeys (*Lagothrix spp.*), as well as all muriquis (*Brachyteles spp.*). These are some of the larger Pan American monkeys that may reach up to 10 kg in body mass, which their prehensile tails can easily handle. Generally, they have complex social structures that have been linked to their morphology (Bjarnason et al., 2015) and cognition (Amici et al., 2008).

We studied three wild Atelid species occurring in the region, *Alouatta palliata* (mantled howler monkeys), *Ateles geoffroyi* (Central American spider monkeys) and *Sapajus imitator* (white-faced capuchin monkeys):

- *A. palliata* (3.1-9 kg): They are some of the most resilient Atelids, and can be found in fragmented habitats (Scheirer et al., 2023) due to their generalist diets (i.e., folivore-frugivore) and low-energy lifestyles (Arroyo-Rodriguez and Duarte Dias, 2010; Jhonson *et al*. 2021). Mantled howlers typically live in multi-male-multi-female groups between 10–15 individuals (Bezanson etc al., 2008). They are categorized as “Vulnerable” by the IUCN (Cortez-Ortiz et al., 2021) and they have a home range of 5-40 ha (Stoner, 1996). Births may occur in any month (Estrada, 1982), and they are often classified as being non-seasonal breeders (Di Bitetti and Janson, 2000), but their reproductive seasonality, if any, may be dependent on population-specific fluctuations in food availability.
- *A. geofroyi* (6-9.4 kg): These spider monkeys live in multi-male-multi-female groups, which separate into smaller, short-term sub-groups depending on several ecological factors (i.e., fission-fusion) (Pinacho-Guendelan et al., 2017). Their group sizes range between 5.6-8 individuals; they use home ranges of 5-962 ha (Asencio et al., 2012). They have a diverse diet and eat fruits, leaves, and insect larvae (González-Zamora et al., 2009). They are currently categorized as being “Endangered” by the IUCN (Cortez-Ortíz et al., 2020). Little is known of the reproductive patterns of these monkeys in the wild, but some studies suggest that there are seasonal variations in male (Hernández-López et al., 2002) and female (Campbell, 2004) reproductive physiology.
- *S. imitator* (2.6-3.9 kg): These capuchin monkeys live in multi-male-multi-female groups that are, on average, made up of 17.2 individuals (Hogan et al., 2019). They have home ranges from 134.4-164 ha and they have an omnivorous diet (Bergstrom et al., 2019). The species is currently considered “Vulnerable” by the IUCN (Williams-Guillén et al., 2021). Reproductively viable females generally give birth every 26.4 months (Melin et al., 2020). Males disperse from their natal group around age four (Jack and Fedigan, 2004). Males are capable of reproduction when they are between 7-10 years of age (Perry et al., 2012), but this may be dependent on their body size and social status (Jack et al., 2014).

### Study Area

The Costa Rican NHP studied live in forest patches of several small-scale private farms in the district of La Rita, Pococi, in the province of Limón, approximately 30 km north of Guapíles, the provincial capital. The community of 100 families, who own government-given lands, are between 2-4 hectares in size. These lands were originally designated as agricultural lands when they were given to local residents in need, and still fulfill this purpose. To promote the conservation of the local biodiversity, the government set in place several regulations that inhibit careless deforestation, especially around creeks and rivers, and any areas set aside by the community as natural reserves. This has resulted in a series of forest patches that follow the many creeks and rivers in the area (Fig. 1).

**Fig. 1:**
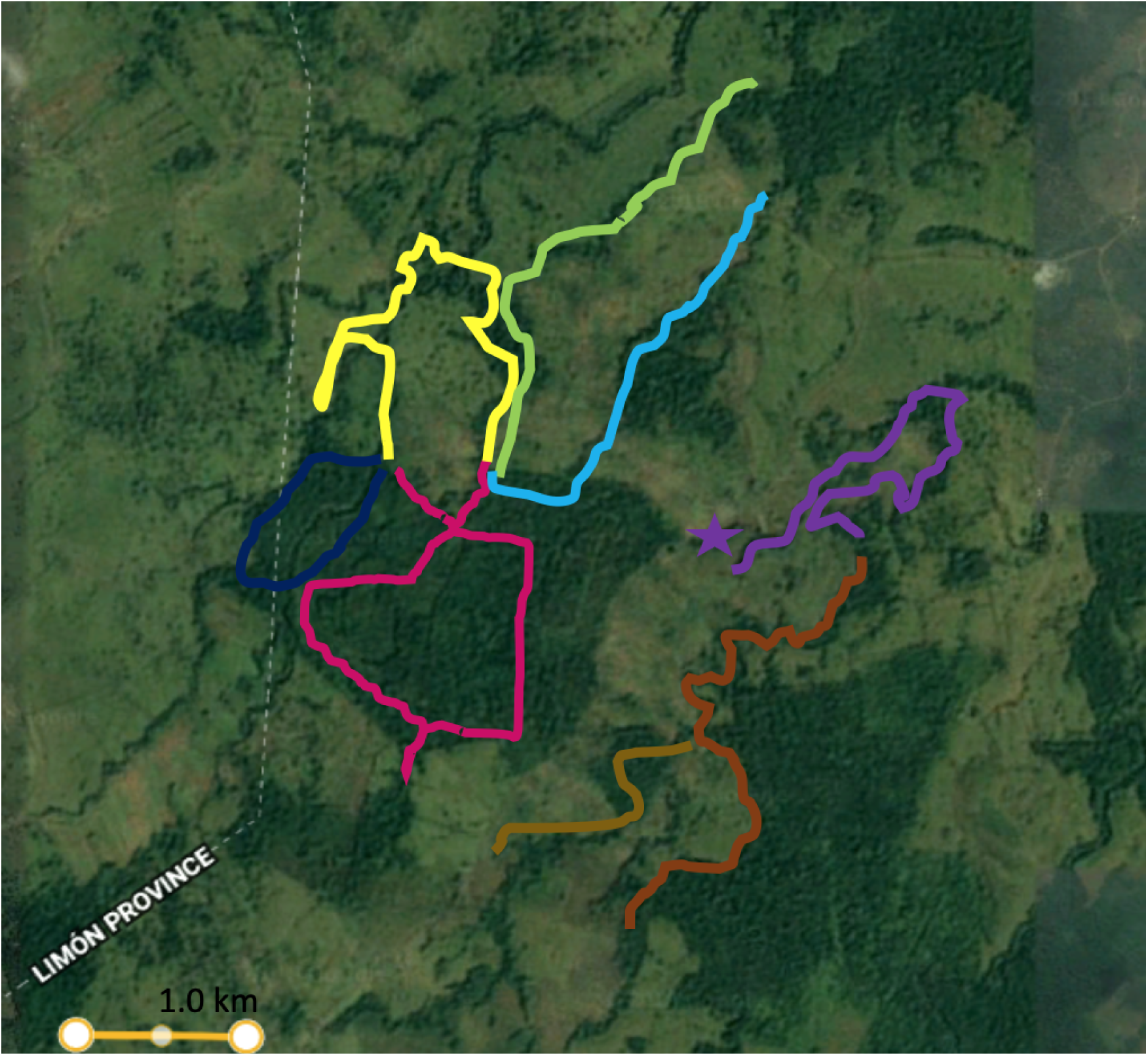
Map of the areas sampled in the district of La Rita in the province of Limón, Costa Rica, Central America. The site is approximately ∼75 kilometers northeast from San Jose, the country’s capital. The 8 forest patches monitored are delineated by the colored lines: Bosque Chonta in dark blue, Rocio-La Teca in red, Rocio-Pedro in yellow, Rocio-Timo in light green, Rocio-Lomas in light blue, La Palma in purple (star shows island), Luis-Esther in dark brown, and Felo-Porvenir is in light brown. Two of the forest patches (i.e., Rocio-Lomas and Felo-Porvenir) contained natural reserves.

Although these forest patches are useful to local wildlife, several species, including the monkeys in this study, are able to use the ground to cross the roads and move between patches, always at an enormous risk. The area is surrounded by large commercial pineapple, palm oil, and banana farms, which employ many of the inhabitants of the settlement. The area is located 1 km north of La Suerte Biological Research Station (LSBRS), a 3-km^2^ tropical rainforest fragment in northeastern Costa Rica (10**°**26’N, 83**°**46’W) (Garber *et al*. 2010). Although LSBRS is protected, forest is being destroyed and fragmented in the surrounding area (Molina, 2015).

The 3 species of wild Atelid monkeys of interest have been studied for decades in Costa Rica, including work at La Selva and adjacent areas (i.e., LSRBS). This work has revealed that *A. paliatta* populations in La Selva are declining and that group sizes are generally bigger in fragmented forests, compared to continuous forests (Bolt et al., 2022), although their behavioral patterns remain the same (Schreier et al., 2021). *A. geofroyi* adapt to fragmented landscape by increasing feeding time, eating less nutrient-rich food items (i.e., leaves) (Chaves et al., 2011), and reducing sub-group size (Rodriguez, 2017). *S. imitator* adapt to fragmented landscapes by socializing in larger groups in smaller home ranges (Tinsley Johnson et al., 2020).

### Data Collection

#### NHP data

Forest patches were sampled 1-3 times monthly, which involved 1-2 trained observers walking the complete transect from end to end, alternating the starting and the ending point. Every time a group of NHP was encountered by the observers, they noted the time of the encounter and remained with the group for at least 30 minutes, and noted the species, group size and number of adults, juveniles, and infants composed the group. The observer also recorded the overall behavior of the group, which was generally foraging, resting, or traveling. *Ad libitum* data was similarly gathered when animals were encountered while we traveled through the patches on non-sampling days or in transit to sample other patches.

#### Climatological data

Historical hourly temperature, humidity, precipitation, rainfall, wind speed, and cloud over data were acquired from OpenWeatherMap. These data were used to characterize the conditions when NHP were encountered.

#### Forest-patch characteristics

The ecology of the 8 forest patches monitored are directly affected by the agricultural practices of the property owners where they are found as such they differ in several general characteristics (Table 1). Each patch has a transect, which we maintain throughout our sampling. Two of the 8 forest patches have a ∼2 km^2^ area designated as a natural reserve where access is limited and the community works together to keep poachers and loggers out with some success. Each reserve has a walkable transect that is maintained by the community for surveillance.

**Table 1:** Route characterizations; the distance of each patch was measured using a handheld GPS. Two of the routes have 2 km2 of natural reserves allocated by the community decades ago.

| Patch ID | Reserve | Distance (km) | General description |
| --- | --- | --- | --- |
| Rocio - La Teca | No | 5.5 | Teak (*Tectona grandis*) production, low biodiversity |
| Rocio - Lomas | Yes | 4 | Includes Las Lomas reserve, lots of water available (muddy soil). High biodiversity |
| Rocio - Timo | No | 4 | Medium stream, high biodiversity, lots of cattle |
| Rocio - Pedro | No | 7 | Large stream, cattle, low biodiversity |
| Luis - Esther | No | 6.5 | Large stream, lots of cattle, medium biodiversity |
| Felo - Porvenir | Yes | 4 | Includes the Sota 2 reserve, high biodiversity |
| Bosque Chonta | No | 4 | Large river traverses it, lots of bromeliads and pejibaye. Occasionally floods. High biodiversity |
| La Palma - Luis | No | 6.5 | Oil palm production, adjacent to a large forest fragment. Medium biodiversity, large forest island present |

### Data Analyses

We used an Information-Theoretic (IT) approach to model the presence of monkeys, independent of species, in each patch, as well as their behavior (i.e., foraging, resting, traveling) depending on 8-9 main effects. This statistical approach facilitates the evaluation of how different models in a given model set (determined *a priori*) fit the data collected, and highlights the explanatory power of one best-fit model or several best-ranked models in the set using Akaike’s Information Criterion corrected for small sample sizes (i.e., AICc) (Burnham and Anderson, 2002; Symonds and Mousalli, 2011). Therefore, the procedure involved calculating several statistical measures from a number of models determined *a priori* (i.e., the model set), based on each model’s AICc value, which is a measure of how well a specific model fits a dataset. In IT terms, AIC approximates the Kullback–Leibler divergence, providing a measure of the relative information loss when using the candidate model to represent the truth.

We calculated differences in AICc between the best-fit model (i.e., model with lowest AICc) and the other models in the set (ΔAICc), the relative likelihood of a model within a given set (AICc Weights), cumulative AICc weights, and the number of times a best-fit (or best-ranked) model was more parsimonious than the lower-ranked model (Evidence Ratios). The ultimate goal of the IT paradigm is to identify the best explanatory variables for the outcome variable being studied/explained. Generally speaking, the best-fit model from the set has the lowest AICc and an AICc Weight larger than 0.90 (Burnham and Anderson, 2002).

We used multi-model averaging for models with cumulative weights up to 0.95 (i.e., 95% best ranked models) for inference making for cases when several best-ranked models were identified (Burnham and Anderson, 2002; Symonds and Mousalli, 2011). Estimates for the parameters in the best-fit models were used to make causal inferences about their effect on the 3 behaviors of interest (i.e., foraging, resting, traveling). In cases where a single ‘best-fit’ model was identified, we used its’ estimates to make inferences about the relationships in the model parameters (e.g., patch name, temp, etc) and our 2 outcome variables modeled (i.e., presence/absence of monkeys, behavioral patterns). If more than one best-ranked models were identified, used model-averaged estimates for each parameter for the 95% best ranked models (i.e., AICc Cumulative Weights < 0.95), which allows us to perform multi-model inferences using model averaging.

All data analyses, summaries, and visualizations were done using Python. Exploratory data analyses were done using the *pandas* library (McKinney 2011) and data visualizations were done using the *seaborn* library (Wazkom 2021).

#### Exploratory Data Analysis

We used 707 instances, 660 when monkeys were encountered and 47 when monkeys were not encountered, to model the presence of the 3 monkey species of interest. To complement our a priori model set, we explored the data collected to understand it and to identify possible confounding factors. We built a 28-model set with models representing the effects on the forest patch ID, the length of the patch, the presence of a reserve, the presence of other monkeys in the patch when sampled, ambient temperature, ambient humidity, rainfall in the hour prior to the encounter, wind speed, and COVID-19 restrictions, as well as some of their interactions.

#### Information-Theoretic Model/Feature Selection

We used 654 encounters to model the foraging, traveling, and resting patterns of the monkeys encountered. We excluded days when no monkeys were found and 6 observations where the behavior was ambiguous. We built a model set with 40 models for the effects of the forest patch ID, the length of the patch, the presence of a reserve, the species go the encountered group, the species of any other groups present in the patch when sampled, group size, ambient temperature, ambient humidity, rainfall in the hour prior to the encounter, wind speed, and some of their interactions.

We built models with the *statsmodel* library (Seabold and Perktold 2010), specifically, we modeled: 1) the presence/absence of monkeys in the 8 forest patches studied with Linear Logistic Regressions algorithms (i.e., *statsmodels.Logit()*); and 2) their behavior during the encounter with Discrete Logistic Regression algorithms (i.e., *statsmodels.LMLogit()*). We pre-processed data using the *sklearn* libraries (Pedregosa et al. 2011) and performed Information-theoretic (i.e., AICc-based) model selection, as well as cross-validation of predictive models, with our custom *douroucoulis* library (somewhat similar to the *AICcmodavg* package in R: Mazerolle 2023).

#### Data pre-processing

Before we fit models we pre-processed the data to meet the specification of each algorithm. This mainly involved coding categorical variables as numerical values. The patch ID was encoded randomly, and the remaining categorical variables were processed using *sklearn*’s *OrdinalEncoder()*.

#### Multi-Model Averaging

We used multi-model averaging for models with cumulative weights up to 0.95 (i.e., 95% best ranked models) for inference making for cases where several best-ranked models were identified (Burnham and Anderson, 2002; Symonds and Mousalli, 2011). We averaged model estimates for each parameter in the best-ranked models to make causal inferences about their effect on the 2 outcome variables of interest. We tested for multicollinearity (Cade, 2015) between the main effects studied with *statsmodels*’ Variance Inflation Factor function (i.e,. *variance_inflation_factor*).

#### Predictive Model Cross-Validation and Hyperparameter tTuning

Once the best explanatory models were identified based on their AICc, ΔAICc, AICc Weights, and Evidence Ratios, we used the best explanatory factors to train 9 classifier Machine Learning Algorithms in order to identify the best performing ones. Once trained, we used k-fold cross-validation to identify the best predictive model using the proportion/percentage of correct predictions over total predictions (i.e., accuracy). For cross-validation, the data were split into two portions: the training set (80%) for developing all base learner models; the test set (20%) to estimate the overall performance of the selected final predictive model.

### Ethical Note

This work is observational and was done non-invasively. We were given permission from the land owners to sample the area after a detailed presentation of our objectives and methods. Our methodology followed the ethical guidelines for fieldwork with wild NHP proposed by the International Primatological Society, the American Society for Primatologists, la Société Francophone de Primatologie, the Primates of Mesoamérica Interest Group, and the Costa Rican government.

## Results

### Exploratory Data Analysis

All three species studied were found in the 8 forest patches monitored (Fig. 2). Monkeys were encountered 660 times over 456 days of the sampling period, which was from October 2018 until August 2020 and from December 2020 until November 2021. Of these 456 sampling days, we did not find monkeys on 47 occasions (10.3%). Of the 660 encounters, the average groups encountered was 1.6 ± 0.4 (mean ± SE) independent of species, and the majority of encounters involved howler monkeys (49%), followed by spider monkeys (21%), and capuchin monkeys (19%).

**Fig. 2:**
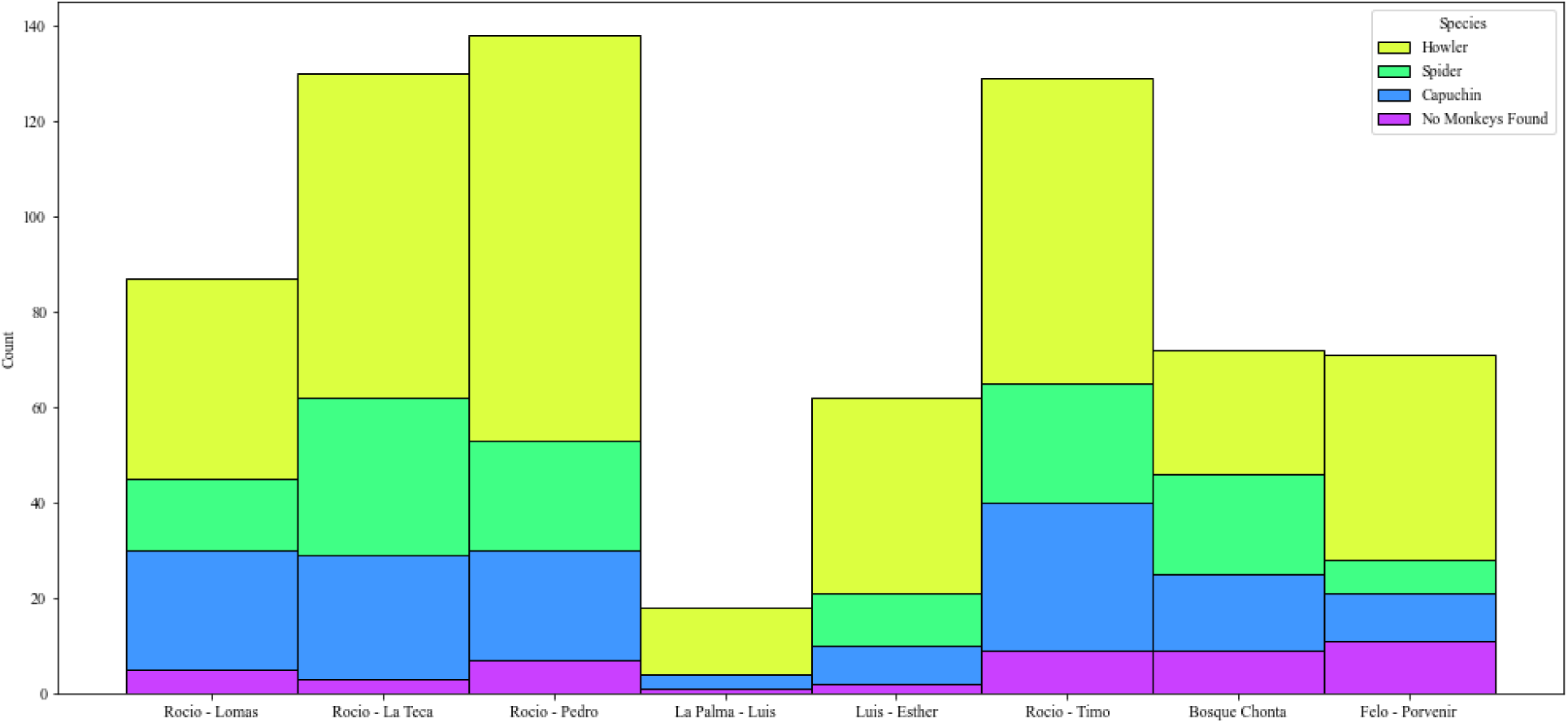
Distribution of the sampling effort for each forest patch. Two of the forest patches (i.e., Rocio-Lomas and Felo-Porvenir) contained natural reserves.

Group sizes were generally similar for capuchins (7.91 individuals ± 0.25; mean ± SEM) and howler monkeys (7.49 ± 0.13 individuals), but smaller for spider monkeys (4.82 individuals ± 0.13) (Fig. 3). Group size generally differed between forest patches; the largest groups were from capuchins encountered in the Felo-Porvenir patch (10.40 individuals ± 1.2; mean ± SEM), whereas the smaller groups were from spider monkeys encountered in the Rocio-La Teca (4.30 individuals ± 0.33).

**Fig. 3:**
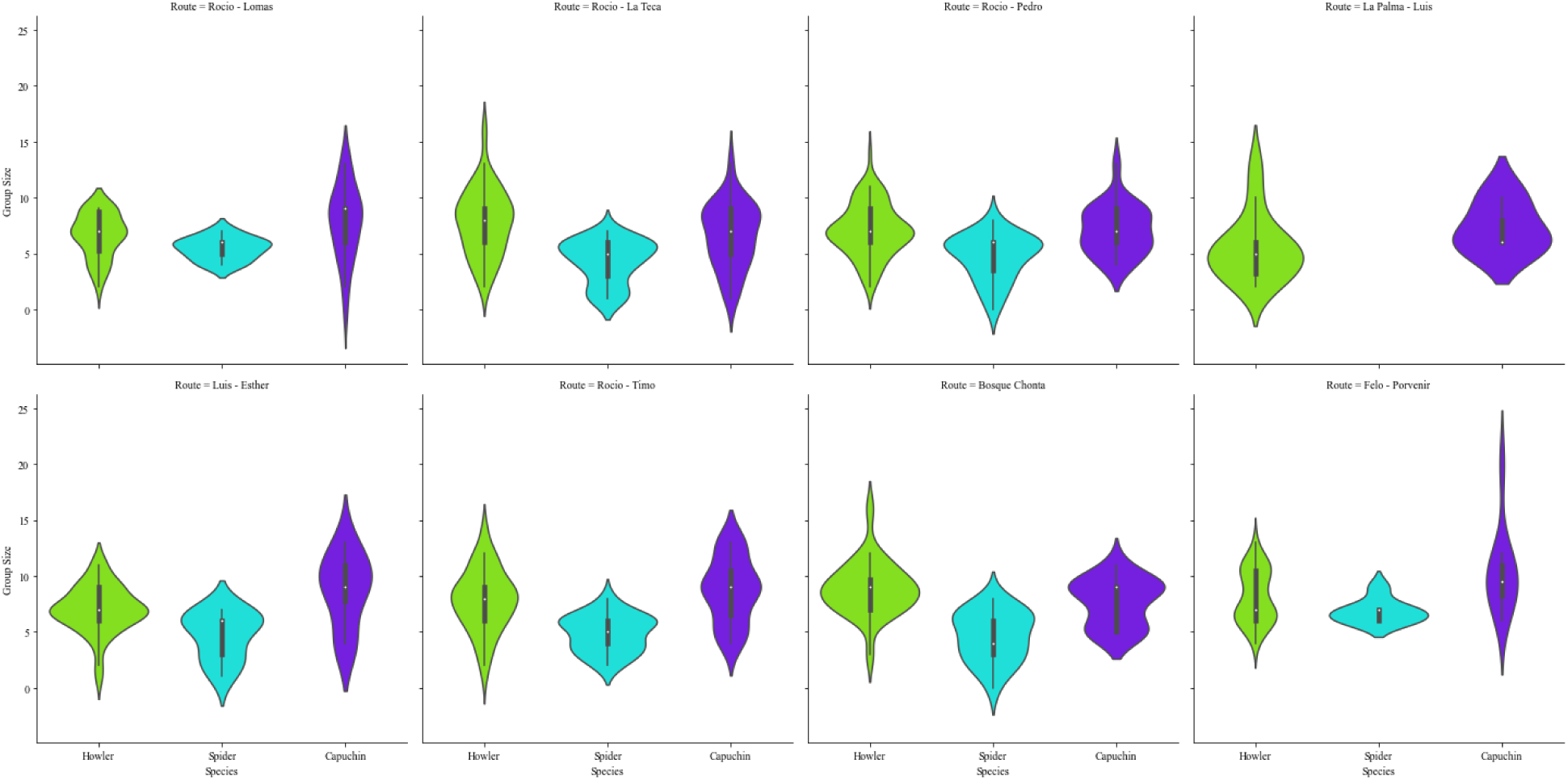
Distribution of group sizes for each species across all forest patches monitored. Two of the forest patches (i.e., Rocio-Lomas and Felo-Porvenir) contained natural reserves.

Behavioral patterns were generally different for each species in each forest patch. Most of the time monkeys were found foraging, followed by traveling, and resting (Fig. 4). Howlers were mainly encountered foraging (235 observations), followed by resting (122 observations) and traveling (23 observations). Spider monkeys were seen mainly traveling (113 observations), foraging (13 observations), and were seen rarely resting (3 observations). Capuchin monkeys were mostly seen traveling (124 observations), foraging (17 observations), and never observed resting (Fig. 4).

**Fig. 4:**
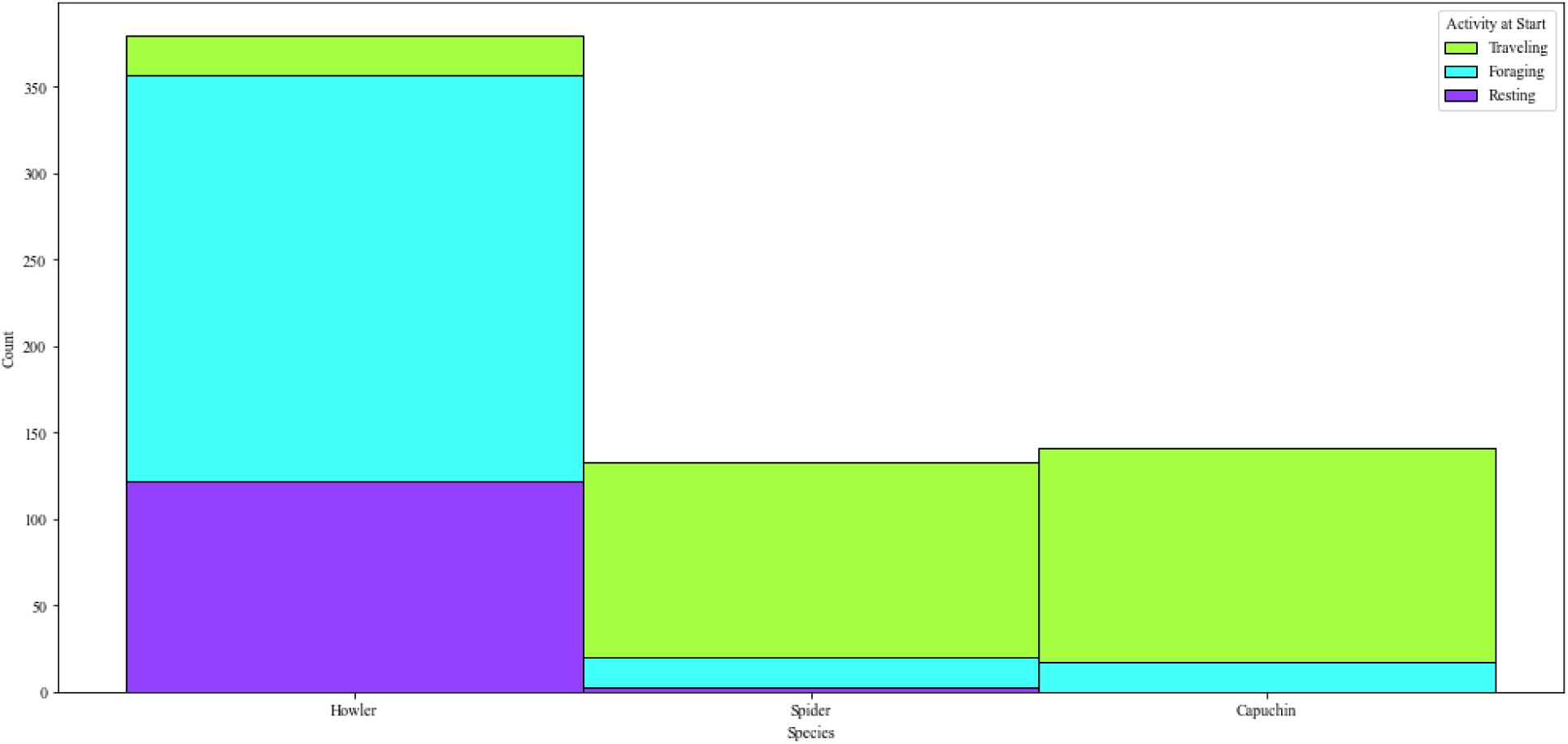
Behavioral patterns for each species.

Climatological conditions fluctuated in a seasonal pattern (Fig. 5). September of 2021 was the warmest (25.83 C± 0.3), most humid month (96.5 % ± 3.0), rainiest (1.14 mm ± 0.33), and windiest (1.81 km/h ± 0.41) month sampled. April of 2019, on the other hand, was the coldest (23.45 C ± 0.05), least humid (88% ± 0.33), driest (0 mm ± 0), and least windy (0.26 km/h ± 0.04) month sampled.

**Fig. 5:**
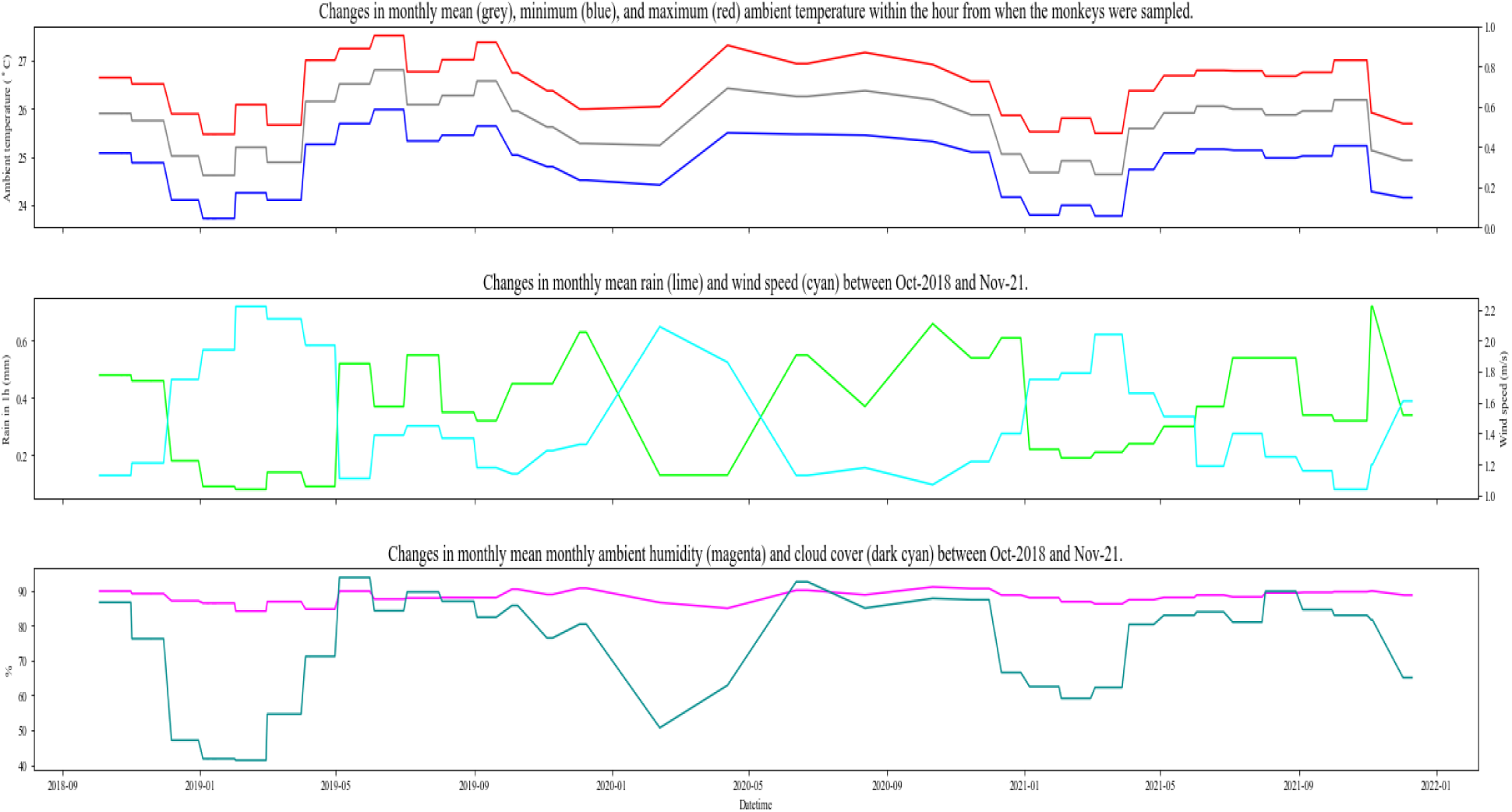
Climatological patterns throughout the sampling period between October 2018 to December 2021.

### Model/Feature Selection

The presence of monkeys in the forest patches monitored was explained by two best-ranked models (Table 2). The first best-ranked model had the presence of other groups in the forest patch and ambient temperature as parameters. The second best-ranked model had the presence of other groups in the forest patch and ambient humidity as parameters. The behavioral patterns of the monkeys studied were explained by 4 best-ranked models with the forest patch ID, the species of monkeys present in the patch when sampled, ambient temperature and humidity, and the group size (Table 2).

**Table 2:** *a posteriori* table showing the best-ranked models explaining the presence and behavior of the monkeys encountered in the 8 forest patches monitored.

| Outcome Variable | Model Name | Parameters | AICc | ΔAICc | AICc Weight | AICc Cumulative Weight | Evidence Ratios |
| --- | --- | --- | --- | --- | --- | --- | --- |
| Presence/<br>Absence | others_temp | Other groups present + temperature | 253.10 | 0 | 0.85 | 0.85 | 1 |
|  | others_hum | Other groups present + humidity | 256.67 | 3.57 | 0.14 | 0.99 | 5.96 |
| Resting,<br>Traveling,<br>Foraging | route_hum_same | Route ID + humidity + species of other group present | 1349.32 | 0 | 0.48 | 0.48 | 1 |
|  | route_temp_same | Route ID + temperature + species present | 1350.53 | 1.21 | 0.26 | 0.75 | 1.83 |
|  | same_hum | Species present + humidity | 1352.66 | 3.34 | 0.09 | 0.84 | 5.36 |
|  | same_groupsize | Species present + group size | 1352.84 | 3.52 | 0.08 | 0.93 | 5.85 |

**Table 3:** Results of k-fold cross-validations for each trained model with the interaction between: the ambient temperature, humidity, and the presence of other groups in the patch, or patch’s name, ambient temperature, humidity, and the presence of other groups in the patch. The accuracy (i.e., proportion of correct predictions over total predictions), thus the closer the value is to 1, the more accurate the model.

| Presence/Absence |  |  | Behavioral Patterns |  |  |
| --- | --- | --- | --- | --- | --- |
| Ambient temp.+Ambient hum.+Other groups present |  |  | Patch+Ambient temp.+Ambient hum.+Other groups present |  |  |
| Trained model | Hyperparameters | Accuracy | Algorithm | Hyperparameters | Accuracy |
| GradientBoostingClassifier |  | 0.96 | AdaBoostClassifier |  | 0.50 |
| LogisticRegression |  | 0.95 | SVC |  | 0.48 |
| SVC |  | 0.95 | XGBoostClassifier |  | 0.48 |
| XGBoostClassifier |  | 0.94 | RandomForestClassifier |  | 0.48 |
| AdaBoostClassifier |  | 0.94 | LogisticRegression |  | 0.45 |
| RandomForestClassifier |  | 0.94 | GradientBoostingClassifier |  | 0.44 |
| KNeighborsClassifier |  | 0.92 | KNeighborsClassifier |  | 0.44 |
| DecisionTreeClassifier |  | 0.90 | GaussianNB |  | 0.44 |
| GaussianNB |  | 0.70 | DecisionTreeClassifier |  | 0.43 |

### Model/Feature-Averaged Estimates of Best-Ranked Models

Estimates for the parameters from the 2 best-ranked models explaining the presence of monkeys in forest patches indicate it was highest in warmer (0.07 ± 0.007; estimate ± SE), more humid (0.02 ± 0.002) days, especially when other groups were seen in the patch (27.99 ± 0.001) (Fig. 6).

**Fig. 6:**
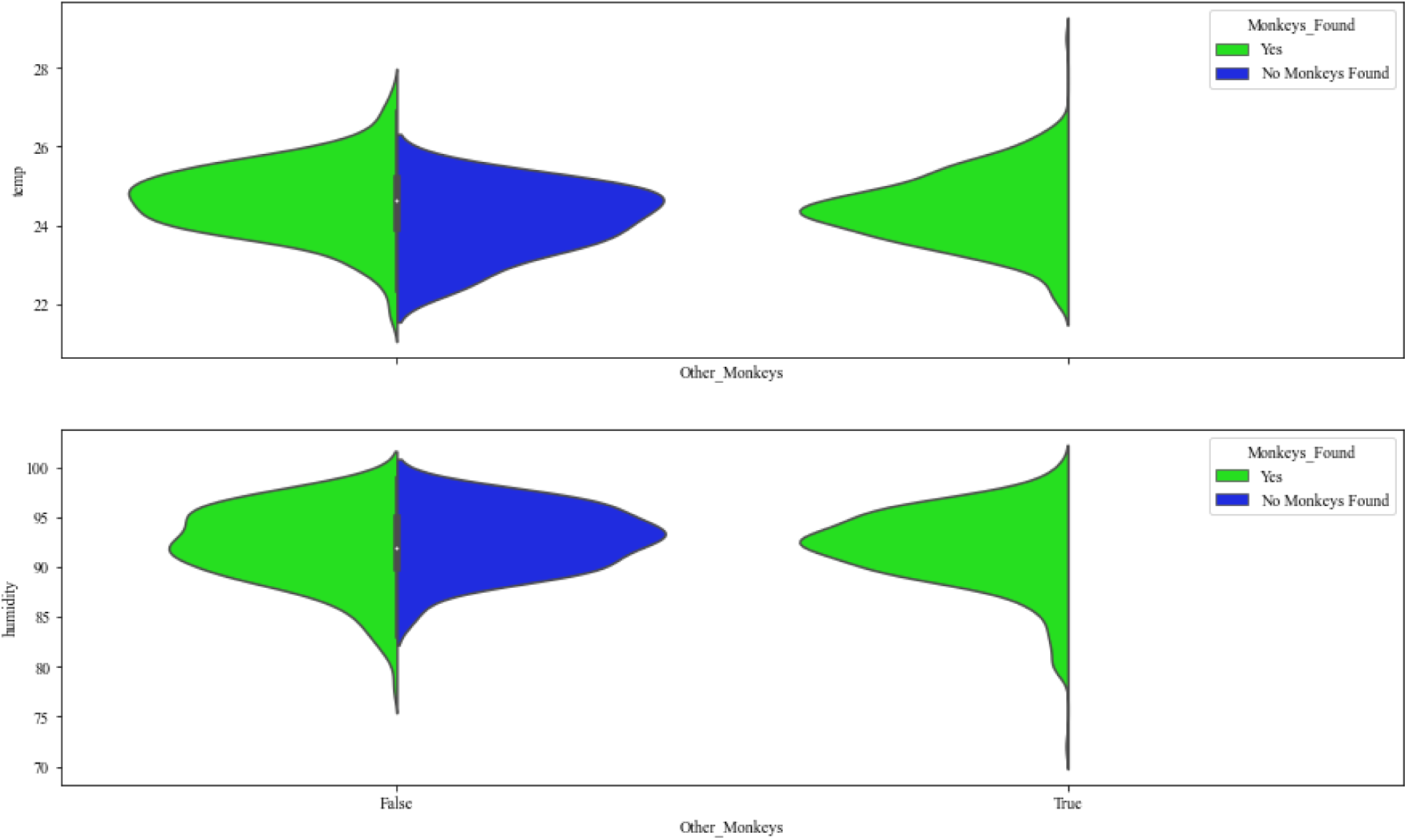
Relationships between the presence/absence of monkeys and ambient temperature and humidity, and the presence of other monkeys in the patch when sampled.

**Fig. 7:**
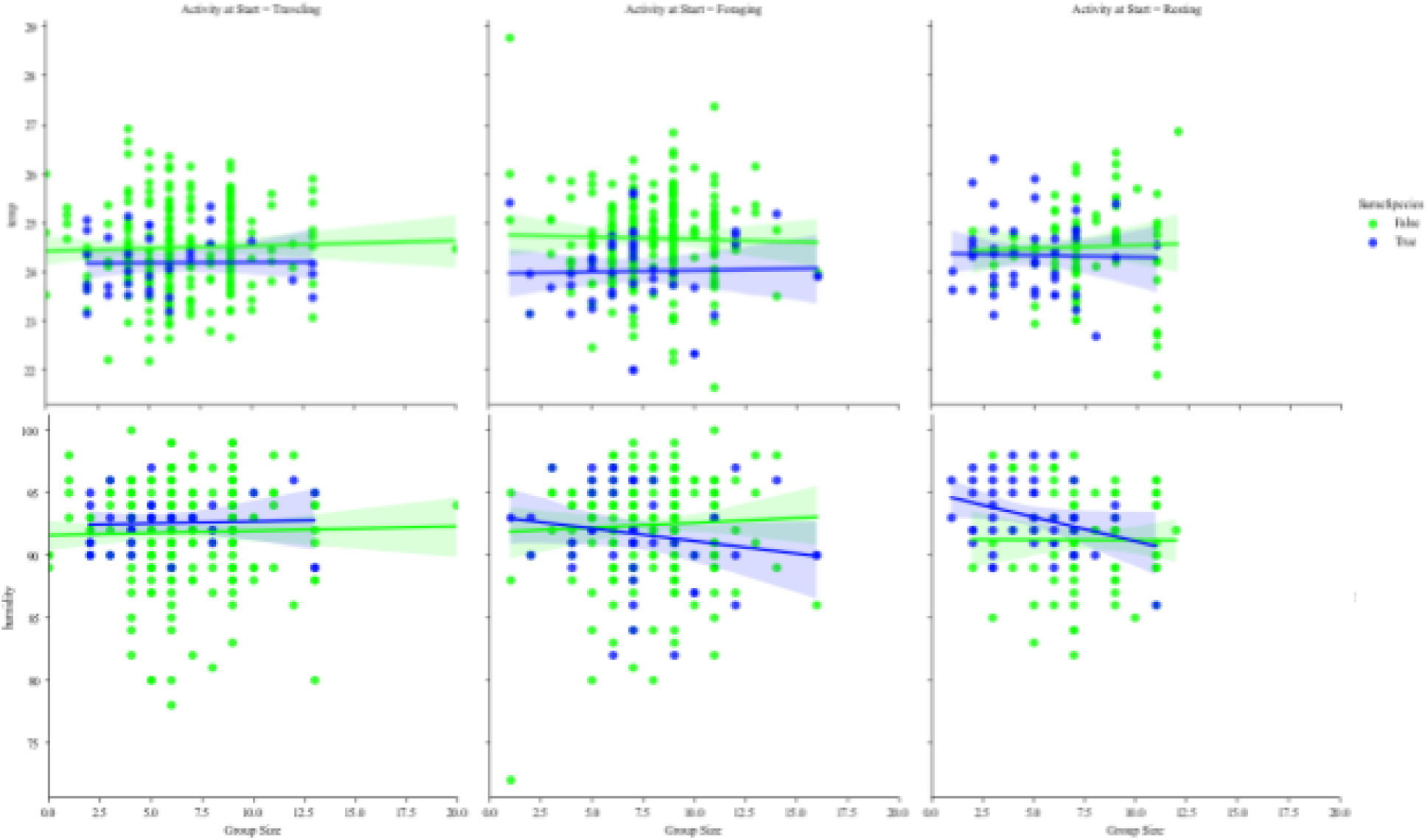
Relationships between the 3 behaviors studied and ambient temperature and humidity, and the species of the other monkeys in the forest patch.

Averages of the estimates for the 4 best explanatory models suggest that, compared to foraging behavior, monkeys were seen more often resting when other groups of the same species were present (0.94 ± 0.25; estimate ± SE); but foraged more often during warmer (−0.06 ± 0.01), and during more humid days (−0.02 ± 0.002). They also foraged more when groups tended to be smaller (−0.14 ± 0.01). The forest patch IDs also explained the variation in our data very well, but since they were (purposely) encoded randomly, it is not possible to determine the relationship from the model estimates (0.12 ± 0.05).

### Cross-Validation of Classifier Algorithms

To validate our best-ranked models, we cross-validated different classifier algorithms with the best-ranked explanatory parameters. For the presence of monkeys in the patches monitored, the Gradient-Boosting Classifier outperformed other algorithms with the highest accuracy of 96%, followed by LogisticRegression (95%). For the behavioral patterns of the monkeys studied, Ada-Boost Classifier and Support-Vector Classifier (SVC) performed best, with an accuracy of 50% and 45%, respectively.

## Discussion

The main threats for the immense biodiversity in Costa Rica and other Central American countries are deforestation and the implementation of non-sustainable agricultural practices. This is because these activities reduce the overall resources available for wildlife to use in their effort to develop, grow, and ultimately reproduce. Non-human primates (NHP), being the most vulnerable vertebrates to extinction, find ways to ameliorate the deleterious effects of habitat loss, adapting to this rapid change (Estrada et al., 2016; Johnson et al., 2023). To understand how NHP deal with this environmental change, we recorded the presence and behavior of 3 Atelid species in a fragmented landscape composed of numerous interconnected forest patches. More specifically, we monitored 8 forest patches with different ecological characteristics (e.g., size, presence of a natural reserve) for the presence/absence and behavior of *Alouatta palliata*, *Ateles geoffroyi*, and *Sapajus imitator.* We hypothesized that monkeys would use the different patches monitored due to their ecological characteristics and the possible benefits they may bring. We used an information-theoretic approach in an attempt to model the multi-dimensionality of the environmental effects studied on the forest patch use by the monkeys studied, which included climatological, demographic, and anthropological factors.

Our main findings showed that the presence of *A. palliata*, *A.geofroyi*, and *S. imitator* in the 8 forest patches studied was best explained by climatological and anthropogenic factors, and not by the forest patch itself. Specifically, monkeys were more likely to visit the forest patches monitored when other monkeys were present and when ambient temperature and humidity were relatively high. In contrast, our analyses of the behavior of the monkeys when they were encountered was explained primarily by the forest patch, independent of size or the presence of a reserve, and by ambient temperature and humidity, the species of other monkey groups encountered in the patch, and their group size. This indicates that the 3 monkeys species studied forest patch use depending climatological, ecological, and anthropological factors.

An insight provided by the results is that the presence of the monkeys in the forest patches was dependent on others being present at the same time, suggesting that any promotion of the presence of one species can lead to the remaining species joining them. This is, in fact, what has been described in other studies where howler monkeys may be the first ones to arrive to disturbed forests, due to their behavioral and physiological characteristics, and other NHP species will arrive later as the forests recuperate. In addition, our analyses of their behavioral patterns suggest that they may be using their daytime activities differently to minimize competition, as foraging was highest when the other groups in the patch were from a different species and they were found resting. This is also corroborated by model estimates of group size that show monkeys, independent of species, tended to forage more often when found in smaller groups, which reduces competition.

One of the important questions remaining to be answered is: what is it about the forest patches studied that may promote a more extensive behavioral repertoire for all of the species studied, and what is it about the reserves (or their management) that is not promoting their usage. For example, there may be a specific nutritional need that is not being fulfilled for *A. geofroyi* and for *S. imitator* because they were mainly seen traveling and foraging. In addition, there may be several microclimatic characteristics that are influencing the usage of each forest patch, as they seem to be ecologically different.

## Conclusion

Taken together, these results suggest that there are several ecological and anthropological factors that explain the usage of a fragmented landscape by *A. palliata*, *A. geofroyi*, and *S. Imitator.* These findings suggest that Atelid monkeys use human-modified landscapes as supplementary food sources and opportunities for traveling (i.e., forest-patch connectivity). This information can be used to develop and implement effective conservation strategies, especially under the paradigm of land-sharing management strategies, and the One Health model of human and NHP well-being.

## Acknowledgements

We would like to thank the community of La Rita for their motivation in conserving the non-human-primates found in their forests, and for their enthusiasm for this study. We would also like to thank all of the participants throughout the years of the Biogas Para Todos efforts, and their interests, in improving the well-being of the community of La Rita. JP would like to give special thanks to Léone Perea-Cruz for the undying support and endless motivation (and smiles).

## Notes

### Competing Interest Statement

The authors have declared no competing interest.

### Summary of Updates

This revision includes the incorporation of predictive models based on the previous exploratory and explanatory predictive models. This was done in an effort to validate the explanatory models from the previous version.

## References

Amici, F., Aureli, F., & Call, J. (2008). Fission-fusion dynamics, behavioral flexibility, and inhibitory control in primates. Current Biology, 18(18), 1415–1419.

Arroyo-Rodríguez, V., & Dias, P. A. D. (2010). Effects of habitat fragmentation and disturbance on howler monkeys: a review. American Journal of Primatology: Official Journal of the American Society of Primatologists, 72(1), 1–16.

Asensio, N., Schaffner, C. M., & Aureli, F. (2012). Variability in core areas of spider monkeys (Ateles geoffroyi) in a tropical dry forest in Costa Rica. Primates, 53, 147–156.

Bergstrom, M. L., Hogan, J. D., Melin, A. D., & Fedigan, L. M. (2019). The nutritional importance of invertebrates to female Cebus capucinus imitator in a highly seasonal tropical dry forest. American journal of physical anthropology, 170(2), 207–216.

Bezanson, M., Garber, P. A., Murphy, J. T., & Premo, L. S. (2008). Patterns of subgrouping and spatial affiliation in a community of mantled howling monkeys (Alouatta palliata). American Journal of Primatology: Official Journal of the American Society of Primatologists, 70(3), 282–293.

Bjarnason, A., Soligo, C., & Elton, S. (2015). Phylogeny, ecology, and morphological evolution in the atelid cranium. International Journal of Primatology, 36, 513–529.

Bolt, L. M., Hadley, C. M., & Schreier, A. L. (2022). Crowded in a fragment: High population density of mantled howler monkeys (Alouatta palliata) in an anthropogenically-disturbed Costa Rican rainforest. Primate Conservation, 36, 1–9.

Burnham K.P., Anderson D.R. (2002) Model selection and multimodel inference: a practical information-theoretic approach. Springer, New York.

Cade, B. S. (2015). Model averaging and muddled multimodel inferences. Ecology, 96(9), 2370–2382.

Campbell, C. J. (2004). Patterns of behavior across reproductive states of free-ranging female black-handed spider monkeys (Ateles geoffroyi). American Journal of Physical Anthropology: The Official Publication of the American Association of Physical Anthropologists, 124(2), 166–176.

Cortes-Ortíz, L.; Rosales-Meda, M.; Williams-Guillén, K.; Solano-Rojas, D.; Méndez-Carvajal, P.G.; de la Torre, S.; Moscoso, P.; Rodríguez, V.; Palacios, E.; Canales-Espinosa, D.; Link, A.; Guzman-Caro, D.; Cornejo, F.M. (2021). “Alouatta palliata”. IUCN Red List of Threatened Species. 2021: e.T39960A190425583. doi:10.2305/IUCN.UK.2021-1.RLTS.T39960A190425583.en

Cortes-Ortíz, L., Solano-Rojas, D., Rosales-Meda, M., Williams-Guillén, K., Méndez-Carvajal, P. G., Marsh, L. K., … & Mittermeier, R. A. (2021). Ateles geoffroyi (Geoffroy’s spider monkey) IUCN Red List Assessment. *See* https://www.iucnredlist.org/species/2279/191688782 *(accessed 10 December 2021)*.

Ceballos, G., Ehrlich, P. R., Pacheco, J., Valverde-Zúñiga, N., & Daily, G. C. (2019). Conservation in human-dominated landscapes: Lessons from the distribution of the Central American squirrel monkey. Biological Conservation, 237, 41–49.

Chaves, Ó. M., Stoner, K. E., & Arroyo-Rodríguez, V. (2011). Seasonal differences in activity patterns of Geoffroy’s spider monkeys (Ateles geoffroyi) living in continuous and fragmented forests in southern Mexico. International Journal of Primatology, 32, 960–973.

Chaves, Ó. M., Morales-Cerdas, V., Calderón-Quirós, J., Azofeifa-Rojas, I., Riba-Hernández, P., Solano-Rojas, D., … & Melin, A. D. (2023). Plant Diversity in the Diet of Costa Rican Primates in Contrasting Habitats: A Meta-Analysis. Diversity, 15(5), 602.

Di Fiore, A., Link, A., & Campbell, C. J. (2010). The atelines: Behavioral and socioecological diversity in a New World radiation. In C. J. Campbell, A. Fuentes, K. C. MacKinnon, S. K. Bearder, & R. Stumpf (Eds.), Primates in perspective (pp. 155–188). Oxford: Oxford University Press.

Estrada, A. (1982). Survey and census of howler monkeys (*Alouatta palliata*) in the rain forest of “Los Tuxtlas,” Veracruz, México. International Journal of Primatology, 2, 363–372.

Estrada, A., Sáenz, J., Muñoz, D., Naranjo, E., Rosales-Meda, M., & Harvey, C. A. (2005). Valor de algunas prácticas agrícolas para la conservación de poblaciones de primates en paisajes fragmentados en mesoamérica. Universidad y Ciencia (México), Número especial 2, páginas 85-94 (2005).

Estrada, A., Raboy, B. E., & Oliveira, L. C. (2012). Agroecosystems and primate conservation in the tropics: a review. American journal of primatology, 74(8), 696–711.

Estrada, A. (2013). Socioeconomic contexts of primate conservation: population, poverty, global economic demands, and sustainable land use. American Journal of Primatology, 75(1), 30–45.

Estrada, A., Garber, P. A., & Chaudhary, A. (2020). Current and future trends in socio-economic, demographic and governance factors affecting global primate conservation. PeerJ, 8, e9816.

Gabriel, D. N. (2013). Habitat use and activity patterns as an indication of fragment quality in a strepsirrhine primate. International Journal of Primatology, 34, 388–406.

Garber, P. A., Molina, A., & Molina, R. L. (2010). Putting the community back in community ecology and education: the role of field schools and private reserves in the ethical training of primatologists. American Journal of Primatology, 72(9), 785–793.

González-Zamora, A., Arroyo-Rodríguez, V., Chaves, Ó. M., Sánchez-López, S., Stoner, K. E., & Riba-Hernández, P. (2009). Diet of spider monkeys (Ateles geoffroyi) in Mesoamérica: current knowledge and future directions. American Journal of Primatology: Official Journal of the American Society of Primatologists, 71(1), 8–20.

Hernández-López, L., Parra, G. C., Cerda-Molina, A. L., Pérez-Bolaños, S. C., Sánchez, V. D., & Mondragón-Ceballos, R. (2002). Sperm quality differences between the rainy and dry seasons in captive black-handed spider monkeys (Ateles geoffroyi). American Journal of Primatology: Official Journal of the American Society of Primatologists, 57(1), 35–41.

Hogan, J. D., Jack, K. M., Campos, F. A., Kalbitzer, U., & Fedigan, L. M. (2019). Group versus population level demographics: An analysis of comparability using long term data on wild white-faced capuchin monkeys (Cebus capucinus imitator). American Journal of Primatology, 81(7), e23027.

Jack, K. M., Sheller, C., & Fedigan, L. M. (2012). Social factors influencing natal dispersal in male white-faced capuchins (*Cebus capucinus*). American journal of primatology, 74(4), 359–365.

Jack, K. M., Schoof, V. A., Sheller, C. R., Rich, C. I., Klingelhofer, P. P., Ziegler, T. E., & Fedigan, L. (2014). Hormonal correlates of male life history stages in wild white-faced capuchin monkeys (Cebus capucinus). General and Comparative Endocrinology, 195, 58–67.

Johnson, C. E., Schreier, A. L., Ramírez, O. V., & Wasserman, M. D. (2023). The Mantled Howler Monkey (Alouatta palliata) Population at La Selva Research Station, Costa Rica: Comparing Censuses in 1992 and 2022. International Journal of Primatology, 1–5.

Johnson, C. E., Tafoya, K. A., Beck, P., Concilio, A., White, K. E., Quirós, R., & Wasserman, M. D. (2023). Primate richness and abundance is driven by both forest structure and conservation scenario in Costa Rica. PLoS One, 18(9), e0290742.

Mazerolle, M. J., Mazerolle, M. M. J. (2017). Package ‘AICcmodavg’. R package, 281.

Melin, A. D., Hogan, J. D., Campos, F. A., Wikberg, E., King-Bailey, G., Webb, S., … & Jack, K. M. (2020). Primate life history, social dynamics, ecology, and conservation: contributions from long-term research in Área de Conservación Guanacaste, Costa Rica. Biotropica, 52(6), 1041–1064.

McKinney W. (2011). Pandas: A foundational Python library for data analysis and statistics. Python for high performance and scientific computing, 14(9): 1–9.

Molina, R. (2015). A Brief History of the Molina Family, and the Birth of Maderas Rainforest Conservancy at the La Suerte and Ometepe Field Stations—A Narrative. Central American Biodiversity: Conservation, Ecology, and a Sustainable Future, 199-213.

Pinacho-Guendulain, B., & Ramos-Fernández, G. (2017). Influence of fruit availability on the fission–fusion dynamics of spider monkeys (Ateles geoffroyi). International Journal of Primatology, 38, 466–484.

Pedregosa F., Varoquaux G., Gramfort A., Michel V., Thirion B., Grisel O., … Duchesnay É. (2011). Scikit-learn: Machine learning in Python. The Journal of machine Learning research, 12: 2825–2830.

Perry, S. (2012). The behavior of wild white-faced capuchins: demography, life history, social relationships, and communication. In Advances in the Study of Behavior (Vol. 44, pp. 135–181). Academic Press.

Rodrigues, M. A. (2017). Female spider monkeys (Ateles geoffroyi) cope with anthropogenic disturbance through fission–fusion dynamics. International journal of primatology, 38, 838–855.

Schreier, A. L., Bolt, L. M., Russell, D. G., Readyhough, T. S., Jacobson, Z. S., Merrigan-Johnson, C., & Coggeshall, E. M. (2021). Mantled howler monkeys (Alouatta palliata) in a Costa Rican forest fragment do not modify activity budgets or spatial cohesion in response to anthropogenic edges. Folia Primatologica, 92(1), 49–57.

Sclater, P. L. (1872). On the Quadrumana found in America north of Panama. Proc. Zool. Soc. London. 1872: 2–8

Seabold, S., & Perktold, J. (2010, June). Statsmodels: Econometric and statistical modeling with python. In Proceedings of the 9th Python in Science Conference (Vol. 57, No. 61, pp. 10-25080).

Stoner, K. E. (1996). Habitat selection and seasonal patterns of activity and foraging of mantled howling monkeys (*Alouatta palliata*) in northeastern Costa Rica. International Journal of Primatology, 17, 1–30.

Symonds, M. R., & Moussalli, A. (2011). A brief guide to model selection, multimodel inference and model averaging in behavioural ecology using Akaike’s information criterion. Behavioral ecology and sociobiology, 65: 13–21.

Tafoya, K. A., Brondizio, E. S., Johnson, C. E., Beck, P., Wallace, M., Quirós, R., & Wasserman, M. D. (2020). Effectiveness of Costa Rica’s conservation portfolio to lower deforestation, protect primates, and increase community participation. Frontiers in Environmental Science, 8, 580724.

Timm, R. M. (1988). A review and reappraisal of the night monkey, *Aotus lemurinus* (Primates: Cebidae), in Costa Rica. Revista de Biología Tropical, 36(2B), 537–540.

Timm, R. M., Wilson, D. E., Clauson, B. L., LaVal, R. K., & Vaughan, C. S. (1989). Mammals of the La Selva–Braulio Carrillo Complex, Costa Rica. North American Fauna.

Tinsley Johnson, E., Benítez, M. E., Fuentes, A., McLean, C. R., Norford, A. B., Ordoñez, J. C., … & Bergman, T. J. (2020). High density of white-faced capuchins (Cebus capucinus) and habitat quality in the Taboga Forest of Costa Rica. American Journal of Primatology, 82(2), e23096.

Waskom M.L. (2021). Seaborn: statistical data visualization. Journal of Open Source Software, 6(60): 3021.

Williams-Guillén, K.; Rosales-Meda, M.; Méndez-Carvajal, P.G.; Solano-Rojas, D.; Urbani, B; Lynch-Alfaro, J.W. (2021). “Cebus imitator”. IUCN Red List of Threatened Species. 2021: e.T81265980A191708420. doi:10.2305/IUCN.UK.2021-1.RLTS.T81265980A191708420.en

Wikberg, E. C., Jack, K. M., Campos, F. A., Bergstrom, M. L., Kawamura, S., & Fedigan, L. M. (2022). Should I stay or should I go now: dispersal decisions and reproductive success in male white-faced capuchins (*Cebus imitator*). Behavioral Ecology and Sociobiology, 76(7), 88.

Zaldívar, M. E., Rocha, O., Glander, K. E., Aguilar, G., Huertas, A. S., Sánchez, R., & Wong, G. (2004). Distribution, ecology, life history, genetic variation, and risk of extinction of nonhuman primates from Costa Rica. Revista de biología tropical, 52(3), 679–693.

